# MOLA: a novel topological data analysis framework for analyzing multiomic loops in precision medicine

**DOI:** 10.64898/2026.07.06.736691

**Authors:** Shael Brown, Bowei Xiao, Kathleen Oros Klein, Jośee Dupuis, Qihuang Zhang, Celia M.T. Greenwood

## Abstract

Multiomic datasets contain complex nonlinear relationships that are often missed by conventional analysis methods. Topological data analysis (TDA) can detect some such patterns (i.e., *loops*), and here we extend previous TDA approaches with MOLA (**M**ulti**O**mic **L**oop **A**nalysis), a framework for multiomic loop visualization, filtering, association with sample-level characteristics and functional characterization. In an Influenza A virus dataset, MOLA better captured anticipated characteristic-dependent epigenetic alterations compared to standard approaches. In a breast cancer cohort, higher loop participation in five genes was associated with increased hazard of progression. MOLA provides an interpretable framework for discovering biologically meaningful multiomic patterns and potential biomarkers.

## 1 Background

Rapid technological advances have made it possible to measure multiple aspects of cell activity and their determinants. For example, genetic sequence in conjunction with epigenetic marks influences transcription, translation, and metabolite production following the central dogma [1], and all these molecular elements can now be measured on high-throughput platforms. However, the functioning of this hierarchy, for any single gene, is highly constrained through the gene’s positioning within a network, where the network system must operate within appropriate boundaries [2, 3]. The availability of rich multi-modal data now allows researchers to search for abnormal patterns in this regulatory network that may be associated with disease.

Nevertheless, identifying multiomic patterns that capture complex regulatory relationships remains a major challenge in precision medicine. A standard approach is to find a low-dimensional representation, of the high-dimensional data, and then to look for patterns in this lower-dimensional latent space. When the goal is to understand heterogeneity as linear combinations of multiomic measurements, matrix methods such as principal component analysis (PCA) and independent component analysis (ICA) can be applied within or across samples. Latent factor methods, such as multiomic factor analysis 2.0 (MOFA+, [4]), can also be applied across samples to identify functionally related genes. However, such linear dimension reductions may miss essential non-linear patterns.

Recently, Gurnari and colleagues [5] proposed a framework for multiomic data analysis that identifies coordinated gene *loop* patterns as a novel class of latent features; we call this framework *Gurnari*. Unlike principal components that can capture global linear variability across a set of genes, *Gurnari* framework-derived patterns are nonlinear and locally structured, capturing relationships within subsets of genes, and may be particularly well suited for periodic biological processes. Even popular nonlinear methods such as UMAP ([6]) are not specifically designed to capture periodic patterns. One example of such a process would be circadian genomics - how patterns of genetic activity and transcription oscillate relative to the 24 hour clock due to biological feedback loops [7]. Because many biological systems can be explained by feedback systems, we expect that latent multiomic loop structures may be important descriptors in many multiomic datasets.

Gurnari et al. [5] identified loop patterns in a multiomic dataset using a tool from topological data analysis (TDA) called harmonic persistence homology [8]. Each loop was represented by a gene weight vector, where weights between zero and one quantified the degree of each gene’s participation in the pattern. Although in [5], the authors visualized these patterns as heatmaps of gene weights, the combination of the weight vectors and the underlying dataset also enabled direct visualization of the loop structure itself; we describe a novel procedure in the Methods section.

Previous applications of the *Gurnari* framework, all described in [5], have followed three steps:

1. Identification of loop patterns and their associated gene weight vectors in each sample of a multiomic dataset.
2. Selection of loops by retaining the top-*k* weight vectors according to a loop-based statistic. However, no principled guidelines for choosing *k* have been established, and the selected value has varied across studies.
3. Downstream analysis of the retained weight vectors using PCA and clustering to identify subtypes of samples.

Our proposed method, **M**ulti**O**mic **L**oop **A**nalysis (MOLA), directly addresses two challenges that arise in the *Gurnari* framework: (a) We provide a principled method for adaptively selecting *k* to improve the robustness and reproducibility of *Gurnari*; and (b) we propose several ways to evaluate the functional characterization of loops, similar to typical enrichment analyses of ranked gene statistics.

In a recent multiomic study of the effects of Influenza A virus (IAV) on macrophages [9], differential signals associated with IAV were identified across multiple omic modalities and genomic loci. We applied MOLA to identify pathways linked to subsets of genes exhibiting loop-like behavior and to quantify differential pathway enrichment across sample characteristics (e.g., IAV status) based on patterns of gene participation in loops. Using MOLA, we recovered expected biological signals, including increased participation of immune, viral, and inflammatory pathways after influenza infection, including the IAV pathway. We also identified differential pathway associations with the ancestry of the study participants. Notably, many of these patterns were not detected using conventional analysis methods. To validate the proposed analytic procedures, we also analyzed multiomic data from breast cancer tumours from The Cancer Genome Atlas (TCGA) [10], and found significant associations between survival and the contribution of certain genes across loops, whereas no significant results were identified in a conventional analytical approach. We then proposed a new approach for characterizing loops based on their functional, rather than gene-level, composition by identifying clusters of functionally-related genes around each loop. Finally, we showed that gene weights, and therefore disease hazard (i.e., instantaneous risk), are sensitive to modifications of individual omic measurements like RNA expression.

## 2 Results

### 2.1 MOLA overview

The steps of the MOLA workflow, applied in both studies, are illustrated in Figure 1 with more details provided in Methods. First, gene-loop structures and their associated gene weight vectors are identified in samples with multiomic data using the *Gurnari* framework. Second, loops are filtered using a statistically-principled inference procedure designed specifically for loops based on a universal null distribution [11], although this filtering step may only be required when samples contain large numbers of loops. Subsequently, when the global multiomic dataset structure is dominated by a single loop, loops (and the samples where they were identified) can be characterized by their clustering of genes in functional pathways around the loop via a “Fourier enrichment” analysis (described in Methods) rather than using individual gene weights. Regardless of the number of retained loops, frameworks of downstream MOLA analyses involve measuring the association between loop-based metrics (e.g. gene weights) and sample-level characteristics (e.g., survival, age, etc.). For genes with significant associations with outcomes we provide an additional downstream example simulation for how to model the sensitivity of the estimated hazard (ratio) to the gene’s RNA expression. The following three subsections illustrate three downstream MOLA analyses on two multiomic datasets.

**Fig. 1:**
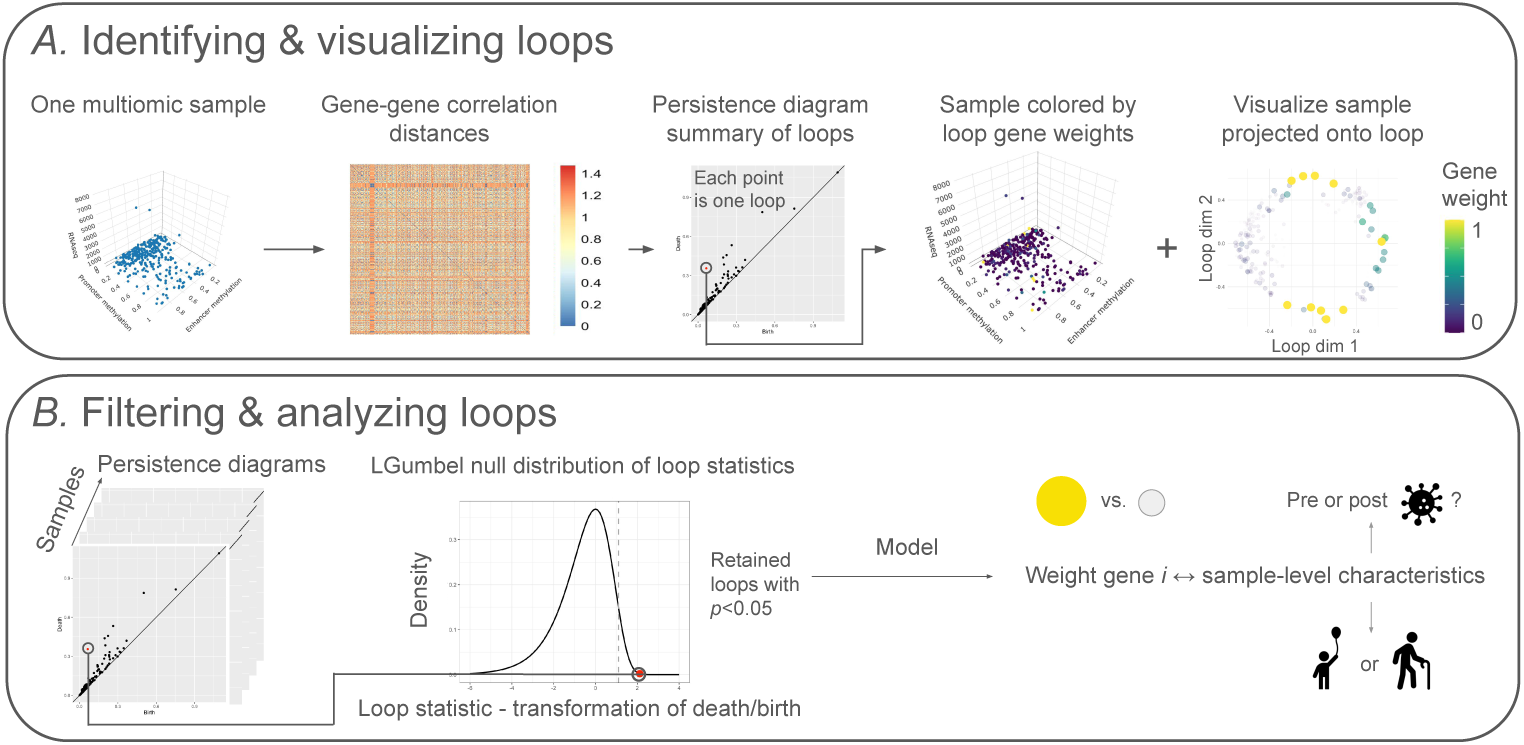
*Steps of the MultiOmic Loop Analysis (MOLA) framework.* (A) Identification and visualization of loop structures. **Left:** a single multiomic sample, projected onto three of the omic dimensions (e.g., RNA-seq, promoter methylation, and enhancer methylation), with each point representing a gene. Using the full dataset (i.e., with all omics), a gene–gene correlation distance matrix is computed (**center-left**) and provided as input to the *Gurnari* framework, which generates persistence diagrams and associated loop weight vectors. In the persistence diagram (**center**), each point represents a loop: the x-axis denotes the birth radius, the correlation distance at which genes connect around the loop circumference, while the y-axis denotes the death radius, the distance at which genes connect across the loop and fill it in. For a highlighted loop, the multiomic sample can be coloured by the loop gene vector (**center-right**), and genes can be projected onto a 2D embedding of the loop (**right**), as described in Methods. (B) Loop filtering procedure and downstream analysis approaches. A transformed loop statistic based on the death-to-birth ratio is computed for all loops across all samples (**left**), the distribution of which follows a standard Gumbel distribution [11] (**left-center**). Only loops with unusually large death/birth ratios, including the highlighted example, are retained for downstream modeling analysis across samples (**right**). We provide two options for such analyses, indicated by the double arrow: evaluating the association between loop gene weights and sample-level characteristics, either by predicting the loop weight of gene *i* according to sample-level characteristics (such as infection status or age), or vice versa.

### 2.2 Study 1: Associations between loop-like epigenetic patterns and sample characteristics in the IAV dataset

We first evaluated our framework on a multiomic dataset of IAV infection in primary human macrophages from 27 healthy male donors of African American or European ancestry [9]. The dataset comprises matched transcriptomic (RNA-seq), and epigenomic (ATAC-seq, whole genome bisulfite sequencing, and four histone marks H3K27ac, H3K4me1, H3K4me3 and H3K27me3) measurements on bulk macrophage cells collected before and 24 hours after infection, yielding 54 paired pre/post-infection samples for downstream analysis. The whole genome omic data of each type were preprocessed to extract signals at gene promoters and their linked enhancers, and to reduce these signals to a single measurement per gene promoter or enhancer, and per datatype (see Methods). Each sample has three associated characteristics (age, ancestry and infection status). We constructed a multiomic data table of *M* genes (rows) by 13 omic measures, where the latter are the summarized signals from 6 epigenetic marks at gene promoters and linked enhancers, plus the gene expression from RNA-seq. We selected *M* = 500 genes with largest variability across samples (see Methods for additional details - this choice was based on computational constraints).

Across the 54 samples, MOLA identified 8323 loops (mean: 154 loops per sample, median: 156 loops per sample), of which 390 (4.7%) were retained based on p-value filtering (see Methods). Gene weights were extracted for each sample’s retained loops. Gene weights were then modeled as a function of sample-level characteristics - age, ancestry, and IAV status, as well as the age-IAV and ancestry-IAV status interactions. The resulting gene-level t-statistics (e.g., the t-values associated with each characteristic for the 500 genes) were then used as input for GO [12], KEGG [13], and Reactome [14] enrichment analyses. The workflow and results of these characteristic-associated enrichment analyses are shown in Figure 2, and the enrichment plots can be found in Supplemental Figures S1-6.

**Fig. 2:**
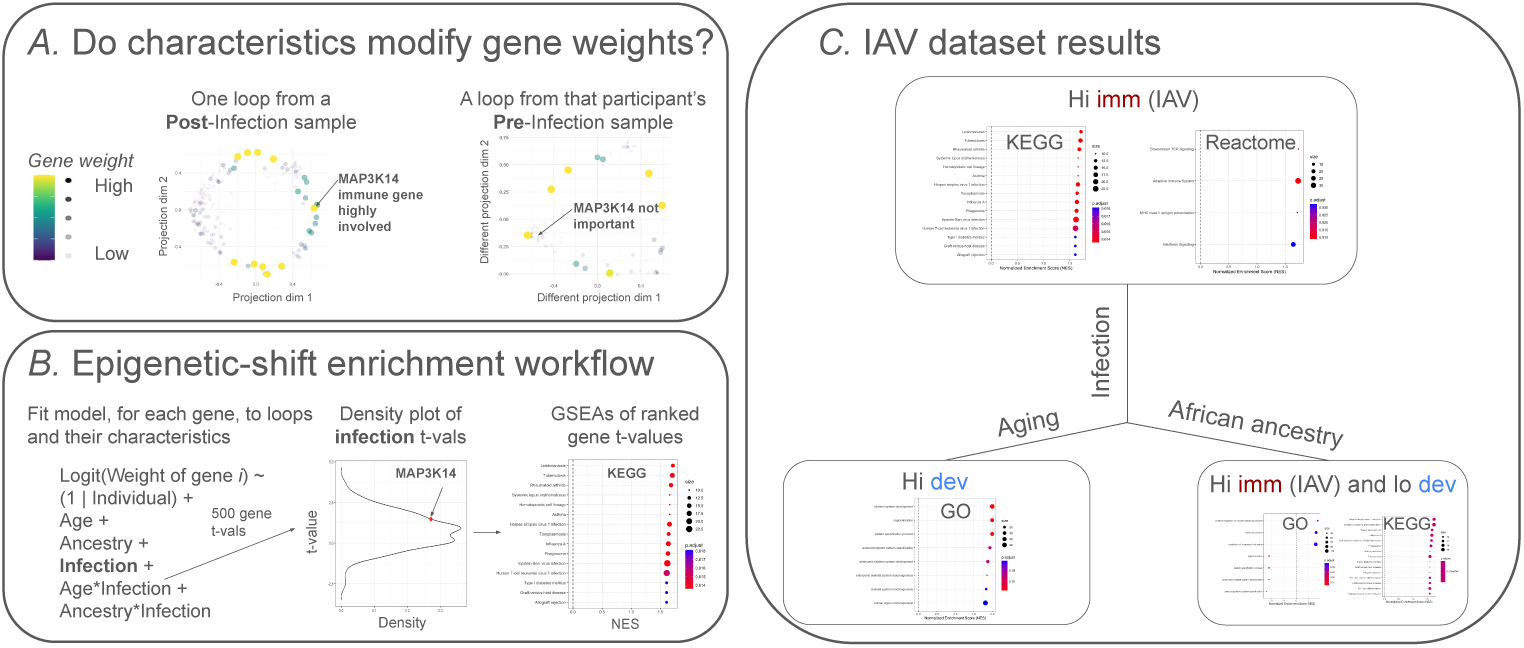
*Characteristic-modified pathway enrichment results in the analysis of the IAV dataset.* (A) Motivation for quantifying characteristic-dependent enrichment patterns. Points are coloured and sized according to weight in the loop. A loop from a post-infection sample (**center**) is shown in a two-dimensional representation, highlighting strong participation of the immune-related gene *MAP3K14*. In contrast, a loop from the same individual’s pre-infection sample (**right**) shows minimal *MAP3K14* participation. This motivates testing whether IAV status (and other sample-level characteristics) increases the participation of immune-related pathways across the 390 retained loops. (B) Analysis workflow. **Left:** for each gene *i*, a mixed-effects model is fit across the 390 retained loops, predicting the logit-transformed gene weight as a function of sample-level characteristics and interactions, with random effects for individual. **Center:** each model term yields a t-value for every gene; the infection-associated t-value for *MAP3K14* is highlighted. **Right:** ranked gene-level t-values are used as input for GSEA analyses (GO, KEGG, and Reactome). The example shown is the KEGG GSEA for infection-associated t-values. (C) Characteristic-associated pathway alterations. Each axis corresponds to a sample-level characteristic with associated GSEA results. Labels indicate whether immune-related (“imm”) or developmental (“dev”) pathways show increased (“Hi”) or decreased (“Lo”) participation in retained loops as the characteristic increases.

Our MOLA results identified two biologically-plausible associations (Figure 2C); infection was associated with increased immune-pathway participation (of genes in retained loops), and aging was associated with increased developmental-pathway participation. An under-representation (i.e., negative enrichment) of developmental pathways were also observed at baseline (not shown in Figure 2, but available in Supplementary Figure S1). Notably, all immune-related enrichment signals involved IAV-associated pathways, supporting the biological validity of the method. Interaction effects between infection and both age and ancestry (not shown in Figure 2, but available in Supplementary Figures S5 & S6) were associated with a reduced immune-pathway participation relative to what might be expected from a model containing only the two individual effects.

These results apply to the selected set of *M* = 500 genes. Repeating the analysis using only the top 300 most variable genes produced different, although partially overlapping, enrichment patterns. More broadly, these findings indicate that the framework is sensitive to the choice of input genes, which should therefore be selected carefully according to the scientific question of interest.

The 500-gene MOLA analysis yielded enrichment patterns distinct from those obtained using two standard alternative approaches: (i) differential analyses performed separately for each omic feature (at promoters and enhancers) with respect to age, ancestry, and infection status, and (ii) enrichment analyses based on Lasso models integrating all 13 omics (some details are provided below, and a more thorough explanation is provided in the Methods).

For (i), generalized linear models (GLMs) were fit to capture association between each sample-level characteristic, such as infection status, age and ancestry, from the 13 separate omic features, and the resulting gene-characteristic test statistics were used as input for Gene Set Enrichment Analysis (GSEA). Enrichment results were mixed; most omics showed enrichment of immune pathways for the infection and ancestry models, but the sign of the effects were inconsistent. Among the 13 models using infection status as the outcome, only the promoter-associated H3K4me1 model identified any enrichment involving the IAV pathway; a signal readily apparent in the MOLA analysis results. Among the 13 omic measures, only the GLMs predicting age from H3K4me3 enhancer signals showed enrichment for developmental pathways; however, unlike MOLA, this enrichment was negative.

For (ii), Lasso models were trained to predict each sample characteristic from 6,500 gene-omic features (500 genes x 13 omic measures). Genes were ranked by the sum of squared coefficients across omic features, and these rankings were used for GSEA; no significant enrichment results were identified from the three models.

Overall, MOLA uniquely recovered strong and consistent enrichment of IAV and other immune-related pathways associated with infection, as well as increased participation of developmental pathways associated with aging.

### 2.3 Study 2: Higher MOLA loop weights for certain genes are associated with reduced survival time among breast cancer patients from TCGA

To evaluate the biological interpretations of MOLA in another dataset, we explored our analytic framework in a multiomic dataset of breast cancer cases from TCGA (BRCA), comprising RNA-seq gene expression, DNA methylation, and copy number variation profiles. After restricting the analysis to female patients with infiltrating ductal or lobular carcinoma, defined menopausal status, and complete survival information (for progression free intervals, as defined in [15] and explained in the Methods), we obtained a curated cohort of 435 samples for integrative analysis. A set of well-known breast-cancer risk- or prognosis-associated clinical characteristics was available for each sample. As in study 1, we selected *M* = 500 highly variable genes for MOLA analysis (rows) with 3 omics measures. Lastly, we removed 28 samples for which there were insufficient computational resources to run the *Gurnari* loop identification step - the remaining 407 samples formed the final cohort.

Out of the total 407 samples, 351 contained one loop, which, when present, typically dominated the sample structure; nearly all remaining samples contained no loops, with three exceptions that contained two loops (but in each of these cases one of the loops was negligible in size and sampling density around the loop, as defined in according to the universal null procedure, and was therefore ignored in subsequent analyses) –therefore, 354 loops were identified, each from its own sample. A mathematical proof of why correlation distances of 3D vectors form a single loop structure is provided in the Methods. Therefore, because each sample is truly characterized by one real loop, the universal null procedure was not appropriate for filtering. This theoretical result can be visualized in the quantile-quantile (QQ) plot for this dataset in Figure 4, where the empirical p-values are much larger than expectation for a uniform distribution. We therefore retained all 354 loops for downstream analyses.

**Fig. 3:**
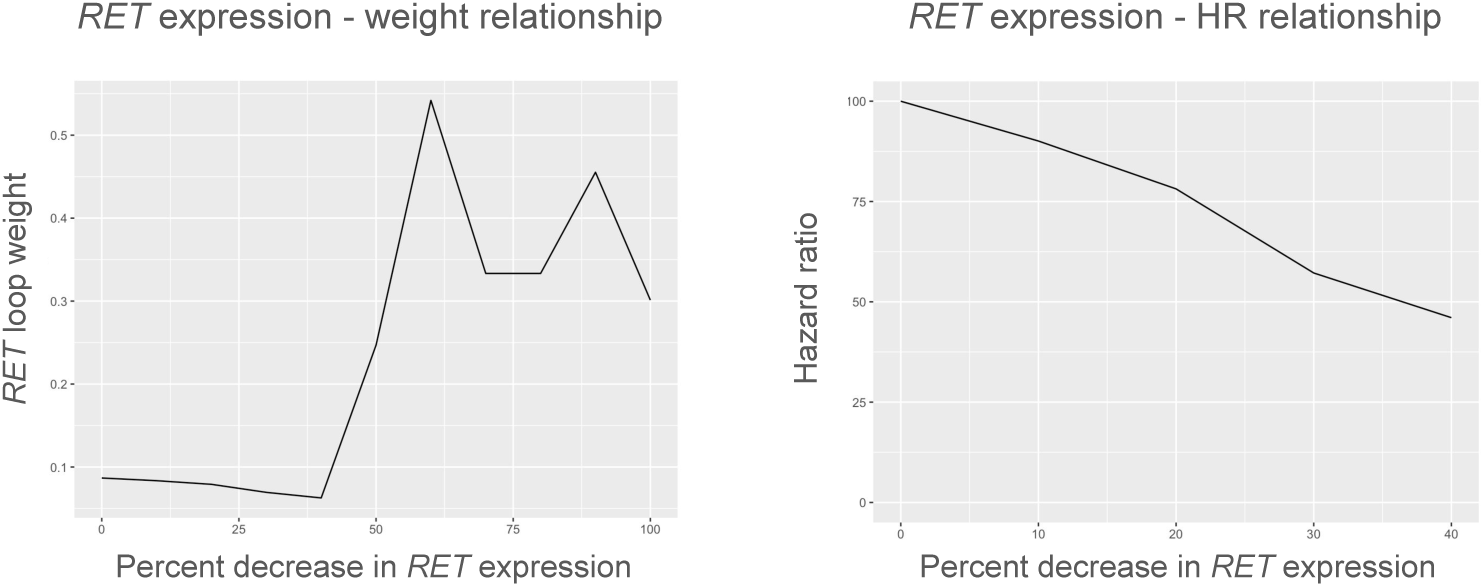
*Sensitivity analysis of RET expression in TCGA sample A7.* **Left:** *RET* loop weight as a function of simulated decreases in *RET* expression. The baseline weight was 0.087 (0% decrease) and declined to approximately 0.063 at a 40% reduction in expression, after which the estimated weights increased sharply. Because these values fell well outside the observed distribution of *RET* weights in the 354 TCGA loops, they were excluded from the hazard ratio analysis. **Right:** Estimated hazard ratio as a function of simulated decreases in *RET* expression (0–40%). The hazard ratio decreased monotonically over this range, reaching a minimum value of 0.46 at a 40% reduction in expression.

**Fig. 4:**
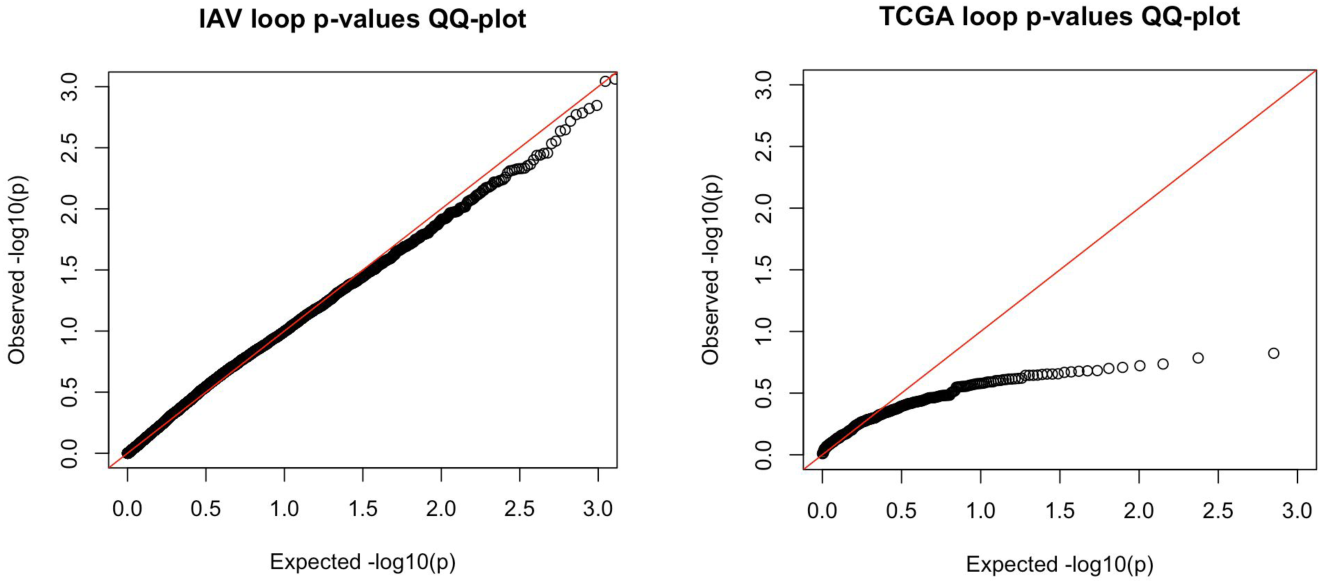
*Loop p-value Q–Q plots for both case studies.* **Left:** The Q–Q plot for the 8323 IAV dataset loop p-values. **Right:** The Q–Q plot for the 354 TCGA dataset loop p-values, after removing the three negligible loops from the three samples that had two loops.

To explore the identified loops, we first examined them based on which pathway’s genes tended to cluster around the loop, i.e. had similar polar coordinates, using a new “Fourier enrichment” framework that we describe in the Methods. A total of 186 loops (53%) were enriched for at least one GO/KEGG term; in those loops an average of roughly 6 terms were enriched. The number of loops that were enriched for a term related to cancer (list of 49 terms provided in Methods) was 87, and 26 loops were enriched specifically for *regulation of apoptotic process*.

Due to the small number of progression events in the final cohort (35), we retained only a small subset of the clinical variables for multivariable modeling. The variables that were consistently significantly related to survival hazard in Cox regression models were (1) pathologic stage, (2) estrogen receptor status and (3) HER2 receptor status, so other variables were excluded from further analysis. For each gene, we used its normalized loop weight in each sample as an extra variable in the Cox model analyzing the 354 loop-containing samples. A false discovery rate (FDR) correction of the 500 gene coefficient p-values yielded 5 genes with significant hazard ratios (Table 1). Some of the genes are known to be implicated in breast cancer and may be suitable targets for future study or intervention - see Table 1.

**Table 1:**
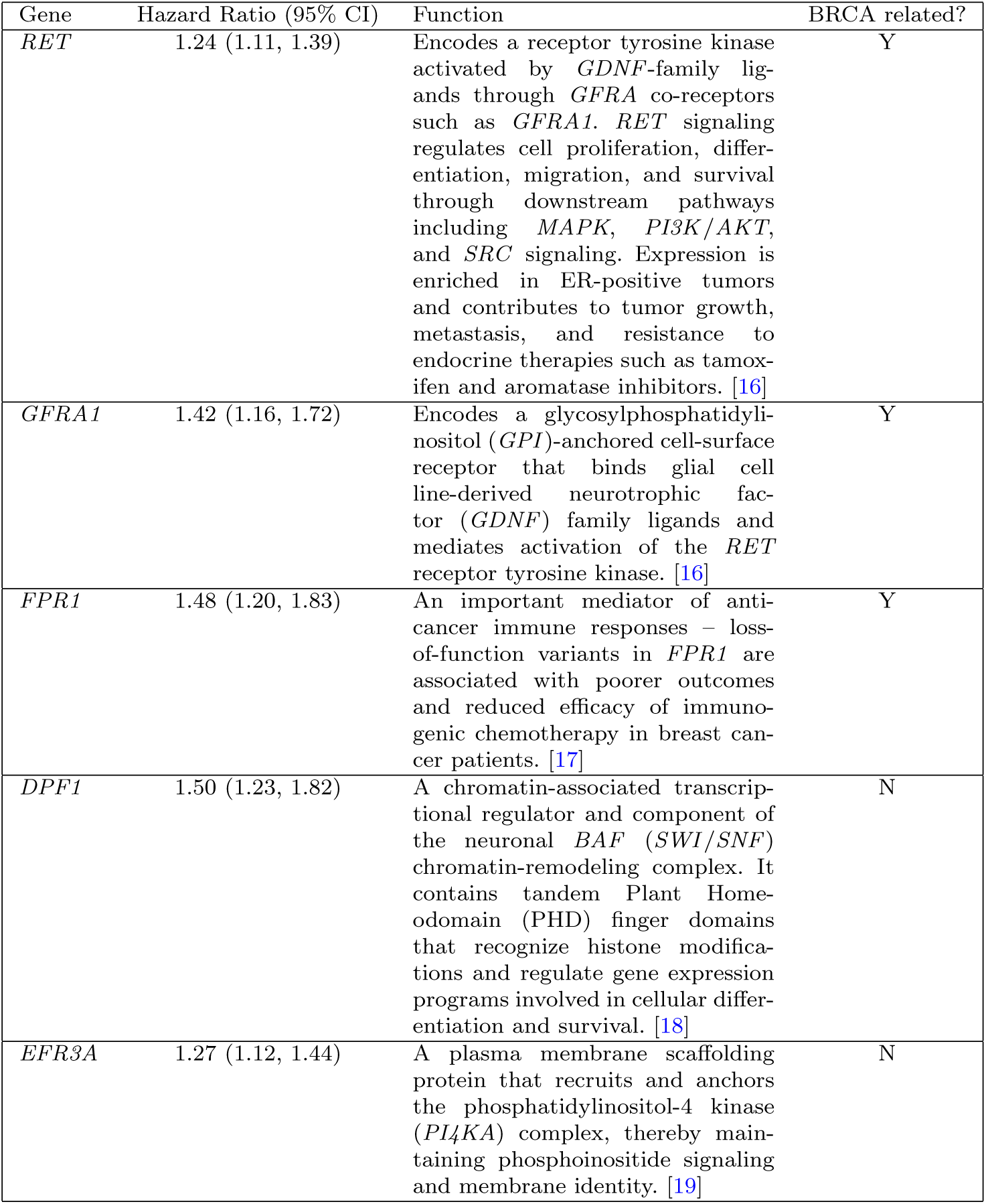
Hazard ratios (HR) and gene functions for five genes with significantly negative prognostic effects in breast cancer survival modeling. 95% confidence interval bounds for the HRs are listed in parentheses next to the estimated value. Whether any strong associations (FDR adjusted p-value *<* 0.05) with breast cancer (”BRCA”) processes are known is noted in the final column.

We then repeated the Cox regression analyses after stratifying samples by histological type (318 infiltrating ductal and 117 infiltrating lobular carcinomas). In the ductal subgroup, only three of the original five genes remained significant (DPF1, FPR1, and EFR3A), while no significant results were detected in the lobular subgroup.

For comparison with a more commonly-used method, we carried out three differential gene analyses, one for each omic measure (RNAseq, CNV and methylation), including each of the *M* = 500 gene’s omic measure in Cox regression models as well as the three retained clinical covariates; no gene coefficients were significant after FDR correction in any of the three models.

### 2.4 Sensitivity of hazard based on individual omic measurements

We next investigated how the estimated hazard for a single TCGA sample (A7-A13E-01, or “A7”) changed in response to alterations in the RNA expression of *RET*, one of the five significant genes identified in our breast cancer analysis. Sample A7 was selected because it experienced a progression event and had the highest *RET* weight among all samples.

We simulated *RET* expression knockdown by reducing its RNA expression by 10%, 20%, …, 90%. For each reduction level *r*, we measured the resulting *RET* weight 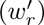 by recalculating the *Gurnari* weights for the modified dataset (i.e., loop) and estimated the corresponding hazard ratio relative to the baseline sample A7, where the *RET* weight was 0.087. Because *RET* weights were standardized before Cox model fitting using their across-loop standard deviation (0.0066), the updated hazard ratio was calculated as the exponential 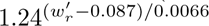 where 1.24 is the estimated hazard ratio associated with a one-standard-deviation increase in *RET* weight. Results are shown in Figure 3.

## 3 Discussion

In this work, we developed MOLA, a novel framework for multiomic analysis that extends the *Gurnari* framework [5]. We demonstrated the utility and broad applicability of MOLA across two studies with completely different goals and designs, and containing different kinds of multiomic data. In the IAV study where the omic data was rich enough to construct separate promoter and enhancer measurements for 6 of the 7 data types, MOLA identified biologically meaningful, sample-characteristic-modified enrichment of loop patterns – including immune response pathways associated with infection and developmental pathways associated with aging – that were not recovered by alternative methods. In the BRCA data from TCGA, MOLA identified five genes whose loop contributions were associated with higher risk of breast cancer progression, demonstrating the clinical potential of our method; no significant results were identified with a standard comparison method. While we did not find statistically-significant associations between survival and other loop-related variables (e.g., including in Cox models one of the loop birth, death or universal null p-value variables and assessing its coefficient for significance; this is discussed further in Methods) such variables may capture novel biological signal in other datasets.

MOLA provides several novel methods for interpreting and analyzing loop features in multiomic datasets. Our loop visualization procedure enables intuitive exploration of many loops. Our loop filtering procedure, based on the universal null framework for TDA, provides a simple heuristic that can be applied consistently across datasets, avoiding the need to tune the *Gurnari* framework’s *k* for each analysis. While stricter multiple-testing correction may be appropriate in some settings, we found it to be unnecessary in the low-dimensional breast cancer study. As a limitation, we note that complex or small loops cannot always be well represented in two-dimensional projections, and may require alternative approaches for interpretation.

We further introduced methods for associating loop-based statistics with sample-level characteristics, including epigenetic alteration and survival analyses. A limitation here is the highly skewed distribution of gene loop weights, which may affect downstream analyses and should be addressed in future work. Finally, we developed Fourier enrichment as an interpretive framework for characterizing loops through pathway-specific gene clustering. The current method detects a single dominant cluster per pathway; future extensions could incorporate the detection of multiple clusters.

MOLA is most effective in datasets that contain both multiomic measurements (ideally, many) and sample-level characteristics. The IAV and TCGA datasets were well suited to the epigenetic alteration enrichment framework introduced in the Influenza case study, whereas the availability of survival outcomes in the breast cancer dataset motivated a separate analysis workflow. More generally, the choice of MOLA analysis framework should be guided by the available data and study objectives.

There is scope to extend the ideas in MOLA in new directions. In the breast cancer dataset where there were only three multiomic measurements per sample, only one loop was identified in the majority of samples. The asymptotic behavior of the significance estimates were unsatisfactory in this context (see Figure 4, right), and alternative strategies could be proposed. Nevertheless, we were still able to obtain meaningful biological insights through Fourier enrichment and associations between gene loop weights and survival outcomes.

By quantitatively linking novel loop patterns to clinically relevant sample-level characteristics, MOLA may enable the discovery of novel biomarkers and biological signals that complement those identified by existing methods. Two of the five significant genes, *RET* and *GFRA1*, from the breast cancer survival analysis are known to be implicated in breast cancer via an overexpression in estrogen-receptor positive breast cancers [16]. Our sensitivity simulation study of the relationship between estimated hazard ratios and *RET* weight in one TCGA sample demonstrates that associations between gene weights and other variables can be highly variable and nonlinear according to modifications in single-omic measurements; future studies could test whether altering the RNA expression of MOLA-identified prognostically-relevant genes through expression knockdown is a viable and effective novel precision medicine strategy.

The general approaches provided in the two case studies would be applicable across a wide range of diseases and multiomic datasets. Beyond precision medicine, MOLA may also be useful in fields such as Neuroscience, where topological approaches to data analysis are increasingly being applied ([20–24]). More broadly, loop visualization has remained a longstanding challenge in topological data analysis with several articles trying to address this problem ([25, 26]), and the wider TDA community may therefore also find value in our approach.

A substantial remaining challenge for the practical application of MOLA is computational cost. The maTilDA package [27] developed for the *Gurnari* framework is highly resource intensive. Computing all loops and gene loadings for samples using *M* = 500 genes required approximately 20 minutes on an Intel Xeon Gold 6226R processor with 1 core and 370 GB RAM, whereas an attempted analysis with *M* = 600 genes required roughly 6 hours on a comparable node with 760 GB RAM. The runtime is dictated by the number of genes, not the dimensionality of the multiomic space, so we recommend restricting analyses to smaller, biologically targeted gene sets aligned with the specific scientific questions under investigation.

The choice of target gene set is likely one of the most critical parameters in the MOLA framework. In our IAV study, results varied depending on whether the top 500 or top 300 highest-variance genes were analyzed. We examined the genes that were ranked 301–500, and they are enriched for immune-related functions, as identified through GO, KEGG, and Reactome over-representation analyses performed with *fgsea* (R package v1.32.4; [28]) using a Benjamini–Hochberg adjusted p-value threshold of 0.05, relative to the full set of 18,187 genes. More broadly, gene set selection can strongly influence downstream enrichment results; for example, restricting analyses to genes already associated with IAV or cancer would almost certainly bias the results toward disease-related pathways. Therefore, care should be taken to select a gene set that is reasonably balanced in composition and not unduly biased.

## 4 Conclusion

MOLA is an interpretable framework for discovering novel multiomic latent patterns that support novel biomarker discovery and risk modeling across precision medicine applications and diverse diseases.

## 5 Methods

### Gene filtering

Empirical testing on our computer systems indicated that *M* = 500 rows was the maximum feasible input size for the MOLA TDA analysis. Accordingly, we applied a variance-based gene filtering procedure to both the IAV and breast cancer datasets.

1. For each omic measure, we computed the variance of every gene across all samples, ignoring missing values.
2. Within each measure, genes were ranked by variance.
3. Each gene was then assigned its highest variance rank across all modalities.
4. The top *M* genes, based on these maximum ranks, were retained for downstream analysis. For most analyses, *M* was selected to be 500; in a sensitivity analysis for the Influenza dataset we also used *M* = 300.

### Identifying multiomic loops

To apply the *Gurnari* framework to define gene loops, we used the accompanying Python package, maTilDA [27]. The framework’s key parameter is a distance function that quantifies dissimilarity between pairs of genes based on their multiomic profiles. Following the original study, we used the Pearson correlation distance, 1 − *ρ*(*g_i_*_1_ *, g_i_*_2_), where *g_i_*_1_ and *g_i_*_2_ denote the omic-value vectors associated with genes *i*_1_ and *i*_2_, respectively, and *ρ*(·, ·) denotes the Pearson correlation. We ran these analyses including either the 500 or 300 most variable genes.

For the Influenza A virus infection analysis, it was necessary to accommodate missing values, a challenge that may also arise in other datasets where MOLA could be applied. We therefore defined a modified distance metric that computes the correlation distance between two gene vectors, *g_i_*_1_ and *g_i_*_2_, using only omic modalities that are non-missing for both genes. In the IAV dataset, each pair of genes in each sample shared measurements for at least 10 of the 13 omic modalities, which we considered sufficient for reliable distance estimation.

For the breast cancer analysis, we filtered in advance to require complete data with no missing values. Our modified distance metric for handling missingness was substantially more computationally expensive than standard correlation distance applied to complete data, and excluding incomplete samples reduced the computational burden associated with processing the larger number of samples. We recommend this approach for other datasets comprised of many samples.

### Visualizing loop patterns

Each loop was identified within a sample-specific gene-by-omic dataset and associated with a weight vector encoding gene participation. To visualize these structures, we applied (point) weighted multidimensional scaling using the R software package vegan version 2.7-3 [29] to the sample’s distance matrix. Constraining the projection by the loop weight vector produced informative two-dimensional visualizations for many loops, examples of which can be seen in Figures 2A and 5-left.

**Fig. 5:**
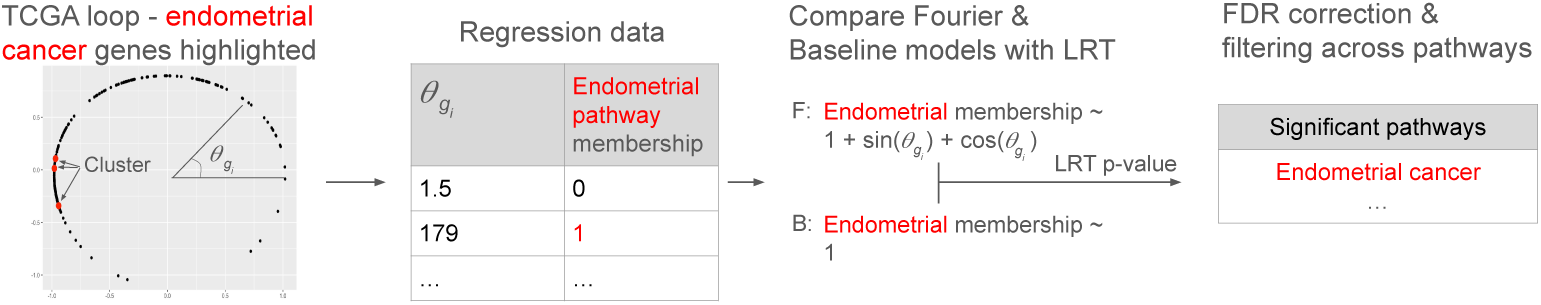
*Fourier enrichment workflow.* **Left:** A TCGA loop in a two-dimensional embedding, where each point represents a gene assigned an angular coordinate (*θ_gi_*). Genes in the endometrial cancer pathway (highlighted in red) cluster near *θ_gi_* = 180^◦^. **Center-left and center-right:** Pathway membership and *θ_gi_* values are used to fit a Fourier model (intercept, cos *θ_gi_*, and sin *θ_gi_* terms) and a baseline intercept-only model. A likelihood ratio test (LRT) compares the models, and resulting pathway p-values. **Right:** Pathway p-values are corrected for multiple testing to identify Fourier-enriched pathways. For this loop, the endometrial cancer pathway is significantly enriched.

### Filtering loops

Loop filtering may be useful in datasets where samples contain many loops, to distinguish loops that are sufficiently well-sampled (according to the universal null distribution) around their circumference from loops that are topological noise [11]. Here we describe the filtering strategy used for the IAV analysis, which may also be applicable to other datasets with similarly high loop counts per sample.

In the original *Gurnari* framework, loop filtering was based on the concept of persistence. In addition to a weight vector, each loop is associated with two scalar quantities: a birth value and a death value [30]. Both correspond to certain special distances, between particular pairs of genes, within the dataset. The birth value is the distance at which genes first form a complete connected cycle around the loop, whereas the death value is the distance at which longer-range connections span the loop and effectively “fill it in.” Persistence is the difference value “death - birth”. Intuitively, the birth value reflects the local sampling density of the loop structure: smaller birth values correspond to denser sampling and more points distributed around the loop. The death value instead reflects a measure of the loop’s size. Persistence therefore quantifies how prominent or stable a loop is across distance distance/connectivity scales.

An alternative, better principled, inferential filtering procedure was proposed by Bobrowski and Skraba [11], in which a certain transformation of death-to-birth ratios across a set of loops was found to follow a parametric null distribution. The transformation aggregates certain information across loops, and outliers are not removed prior to doing so. This point was addressed in [11], which argued that unless there is an unusually large number of real loops in a dataset the transformation will still follow the parametric null distribution. Under this framework, the null hypothesis for a given loop is that its ratio does not exceed the value expected under the null distribution, implying that the loop reflects noise rather than signal. In the original paper, it was recommended to apply a filter to *L* loops based on thresholds defined by a multiple testing correction such as the Bonferroni applied to *L* tests.

There are two ways this procedure can be applied: independently within each sample (i.e., filtering loops separately for each sample) or jointly across all samples by pooling all loops before estimating the loop statistic transformation parameters. Although the sample-wise approach is simpler, the pooled approach provides a more robust estimate of the null distribution across the full cohort. For example, a loop that appears significant within a single sample may not appear significant when considered in the context of all samples, a distinction captured only by the pooled analysis.

Compared with the workflow in [5], our approach provides a more systematic and probabilistically motivated filtering strategy:

- For the IAV analysis, we applied the inferential null procedure to all loops from all samples simultaneously. Application of a Bonferroni threshold for the number of loops in total did not lead to retention of any loops. Therefore, we applied a filter using *p <* 0.05 to select loops for further exploration. We note that biologically meaningful enrichment patterns, identified when applying this *p*-value threshold, were lost when no filtering was applied and all loops were analyzed.
- Most samples from the breast cancer dataset contained at most one loop, all of which were retained for further characterization. Three samples contained a second negligible loop which were ignored.

The Q–Q plots for both analyses are shown in Figure 4.

The predominance of a single loop in the TCGA multiomic samples admits a simple mathematical explanation. Let ℝ^3^ denote the space of length-3 vectors, and let **x** = (*x*_1_*, x*_2_*, x*_3_) and **y** = (*y*_1_*, y*_2_*, y*_3_) be two such vectors (treated as fixed, not random variables). Additionally, we define (Pearson) correlation distance as *d*_cor_(**x**, **y**) = 1 − *ρ*^(**x**, **y**).

Since correlation is translation invariant, distances remain unaffected when modifying a vector by subtracting its third coordinate (treated as a scalar constant) along each dimension: *d*_cor_(**x**, **y**) = *d*_cor_((*x*_1_ − *x*_3_*, x*_2_ − *x*_3_, 0), **y**). Therefore, we can assume that distances are computed in the 2-dimensional plane (ignoring the third coordinate), i.e. **x** = (*x*_1_*, x*_2_, 0) and **y** = (*y*_1_*, y*_2_, 0).

Similarly, since correlation is scale invariant, distances remain unaffected when modifying (i.e., multiplying by a positive scalar) the length of a vector. One example of such a modification is dividing a vector by its length: 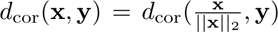, where 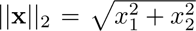 is the Euclidean norm. So, we can assume that all vectors have norm 1, i.e. that 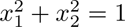.

Therefore, the structure induced on R^3^ by correlation distances projects all data points (genes, in the case of MOLA) onto a unit circle.

### Modeling epigenetic alterations in enrichment patterns across loops in the Influenza A dataset

One of the original applications in [5] compared STRING DB [31] enrichment results between two cohorts (pre- and post-treatment) qualitatively, whereas the goal of MOLA is to enable such comparisons quantitatively. As motivation, Figure 2A illustrates that participation of the immune-related gene *MAP3K14* is high in a particular loop from a post-infection sample in the Influenza A virus infection dataset, but low in a pre-infection sample from the same individual. This naturally raises the question of whether *MAP3K14*, other immune-related genes, and related functional pathways exhibit differential loop participation according to infection status, across the 390 retained loops.

For the Influenza A virus infection analysis, we indexed the 500 genes by *g_i_*, 1 ≤ *i* ≤ 500, and the 390 retained loops by *l*, 1 ≤ *l* ≤ 390. Because each loop originated from a sample with associated characteristic data, we defined the following variables for each loop *l*:

- Age*_l_* - the age of the individual corresponding to loop *l*.
- Ancestry*_l_* - ancestry indicator for the corresponding sample, coded as 1 for African American ancestry and 0 for European American ancestry.
- Infection*_l_* - infection-status indicator for the corresponding sample, coded as 1 for post-infection and 0 for pre-infection.
- Individual*_l_* - participant identifier corresponding to loop *l*. Because each participant contributed both pre- and post-infection samples, this variable accounts for repeated measurements.

Accounting for pre-post pairings in the study, we added an individual level random effect to the models for Individual*_l_*, and fit 500 mixed-effect linear models, one for each gene, of the form:

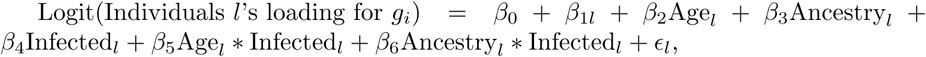

where 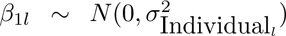 and 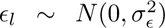. Each model was fit with *l* = 1*, . . .* 390 data points, ranging over the 390 retained loops.

The primary outputs of this stage of the analysis were the model t-statistics. For example, the age characteristic produced one t-value for each gene *g_i_*, yielding a total of 500 gene-specific statistics. For each transformation, the resulting vector of 500 gene t-values was used as input for three gene set enrichment analyses (GSEAs): GO, KEGG and Reactome. Enrichment results for each characteristic were then compared across all three GSEAs to identify consistent biological signals.

Gene set enrichment analysis (GSEA) was performed using the R packages clusterProfiler version 4.14.6 [32] and ReactomePA version 1.50.0 [33], in R version 4.4.2. The 500 genes were ranked according to t-statistics, mapped to Entrez Gene identifiers using the org.Hs.eg.db annotation database R package version 3.20.0 [34], and analyzed using a pre-ranked GSEA approach. Enrichment analysis was conducted against GO, KEGG, and Reactome pathway databases. Gene sets containing fewer than 10 genes or more than 500 genes were excluded. Enrichment scores were calculated using a permutation-based approach (1000 permutations), and statistical significance was assessed using Benjamini–Hochberg false discovery rate (FDR) correction. Pathways with FDR-adjusted p-value *<* 0.05 were considered significant. Normalized enrichment scores (NES), adjusted p-values, and leading-edge subsets were computed using default clusterProfiler and ReactomePA parameters. For a GSEA of the t-values for a characteristic (such as age), positive NES indicates enrichment in upregulated genes (e.g., genes that have higher loop loading/contribution in the cluster loops for samples with larger Age), while negative NES indicates enrichment in downregulated genes (e.g., genes that have lower loop loading/contribution in the cluster loops for samples with larger Age).

### Comparison frameworks for epigenetic-alterations analysis

We compared our new epigenetic alteration framework for loop patterns against two standard analytical approaches.

In the first approach, gene-level associations with characteristics were estimated using a differential expression framework applied independently to each sample-level characteristic – age, infection and ancestry.

For count-based omics data, differential analyses for age, ancestry and infection status were performed separately using DESeq2 [35] in R. A generalized linear model was then fit, predicting the omic measure from each sample-level characteristic. Statistical significance for the coefficients were assessed using Wald tests, yielding signed test statistics for each omic measure. For continuous-valued omics data, a similarly-structured differential analysis was performed using the limma package [36], with empirical Bayes moderation applied to improve variance estimation. In either case, gene t-statistics corresponding to the sample-level characteristics were extracted as feature-level association scores.

The resulting ranked vector of signed test statistics, across genes for each sample-level characteristic, represents the relationship between omic features and characteristics, and was input into 13 separate GO, KEGG and Reactome GSEAs (one for each omic measure) to identify pathway-level trends.

In the second approach, gene-level associations with characteristics were estimated as aggregated (sum of squared) coefficients of Lasso models, predicting sample-level characteristics from gene-omics (of which there are 500 genes x 13 omics, for a total of 6,500 features). Features were median-imputed (i.e., 17 features with partial missingness). There were no missing values for 6,301 features, and 182 features with all missing values were removed. All retained features were then normalized. For categorical or binary characteristics (i.e., infection and ancestry), logistic elastic-net regression was used; for continuous characteristics (i.e., age), Gaussian elastic-net regression was used. The elastic-net mixing parameter *α* was selected by 5-fold cross-validation across the 54 samples over a grid of *α* values, and the final model was fit using the cross-validated regularization parameter. This procedure identifies sparse, multivariate combinations of geneomic features that jointly model the characteristics while accounting for correlations.

Feature-level regression coefficients were then aggregated to the gene level; for each gene, coefficients across all associated omic features were summarized using the Euclidean norm, yielding a non-negative gene-level statistic that captures the overall strength of multivariable association between that gene’s omic profile and sample characteristics. These gene statistics were then input into GO, KEGG and Reactome GSEAs for each sample-level characteristic.

### Fourier enrichment analysis of loops

A functional characterization of loops can either indicate (i) whether genes from particular pathways tend to participate highly in the loop, or (ii) whether genes from particular pathways tend to cluster, by their polar coordinates, around the loop. We did not find evidence that (i) would be a meaningful representation – no enrichment pathways of raw gene loop weights were identified in any of the IAV or breast cancer retained loops.

Typical enrichment analyses require an ordered list of gene statistics with a defined “top” and “bottom,” making them unsuitable for periodic statistics that vary around a loop. To identify pathways whose genes cluster *anywhere* along a loop, we developed a novel method termed “Fourier enrichment analysis.” This approach is appropriate only when the loop structure is well defined and can be characterized by a single latent angular variable that uniquely determines gene positions along the loop. In our breast cancer study, this condition was satisfied. Using a gene–pathway binary membership matrix, we found that several biologically relevant pathways cluster according to the loop angle variable in our breast cancer sample loops. The steps of our Fourier enrichment procedure are shown in Figure 5.

A loop’s embedding (based on the visualization procedure) admits gene angles (*θ_gi_*), and for each enrichment pathway/term there is a binary membership vector of participating genes. This data is input to a “Fourier” linear regression model, Membership ∼ *β*_0_ + *β*_1_ sin(*θ_gi_*) + *β*_2_ cos(*θ_gi_*), and a baseline model, Membership ∼ *β*_0_. A likelihood ratio test (LRT, for testing the hypothesis that *β*_1_ = *β*_2_ = 0) produces a p-value - low values are indicative of a strong cluster of member-genes somewhere around the loop. The collection of p-values, across pathways/terms, for the loop are then adjusted for multiple comparisons.

Out of 460 unique pathways that were enriched in at least one of the 354 loops, 49 were highly related to various themes of cancer processes, and these themes & terms can be seen in Table 2

**Table 2:** Cancer-associated enrichment terms.

| **Cancer-related theme** | **Enriched terms** |
| --- | --- |
| Canonical cancer pathways | Bladder cancer; Breast cancer; Endometrial cancer; Gastric cancer; Hepatocellular carcinoma; Melanoma; Chronic myeloid leukemia |
| Carcinogenesis and DNA damage | Chemical carcinogenesis – DNA adducts; Chemical carcinogenesis – reactive oxygen species; Viral carcinogenesis; DNA damage response; p53 signaling pathway |
| Oncogenic signaling | Wnt signaling pathway; Positive regulation of canonical Wnt signaling pathway; ERK1 and ERK2 cascade; Positive regulation of ERK1 and ERK2 cascade; Fibroblast growth factor receptor signaling pathway; Notch signaling pathway; NF-κB transcription factor activity |
| Cell proliferation and cell-cycle control | Cell division; Regulation of cell cycle; Positive regulation of cell division; Positive regulation of cell growth; Regulation of cell population proliferation; Positive regulation of stem cell proliferation |
| Tumor invasion, adhesion, and angiogenesis | Cell migration; Regulation of cell adhesion; Integrin-mediated signaling pathway; Positive regulation of angiogenesis; Positive regulation of sprouting angiogenesis |
| Cancer-associated gene regulation | Regulation of DNA-templated transcription; Regulation of gene expression; Positive regulation of transcription elongation by RNA polymerase II; MicroRNAs in cancer; Transcriptional misregulation in cancer |
| Tumor immune and viral oncology | PD-L1 expression and PD-1 checkpoint pathway in cancer; Human T-cell leukemia virus 1 infection; Viral carcinogenesis |
| Cancer metabolism | Choline metabolism in cancer |

### IAV dataset

Raw FASTQ files were available for macrophage samples from individuals before and after *in vitro* infection. However, analyses were restricted to the 27 (out of 39) individuals with complete multi-omic measurements of all 7 types. To ensure comparability across epigenetic data types, all FASTQ reads were re-aligned to the GRCh38 v87 reference genome using data-type-specific aligners: Bismark v0.18.1 for methylation data, BWA v0.7.17 for ATAC-seq and ChIP-seq data, and STAR v2.7.8a together with HTSeq v2.0.4 for RNA-seq data. Data-type-specific preprocessing and quality control were performed using GenPipes v6.0.0 with default parameter settings [37].

For ATAC-seq and ChIP-seq data, peaks were called using MACS2 to generate genome-wide peak sets for downstream analyses. Narrow peaks were identified for ATAC-seq, H3K27ac, H3K4me1, and H3K4me3, whereas broad peaks were identified for H3K27me3. For bisulfite sequencing data, methylated read counts and total read coverage were quantified at predefined CpG sites using Bismark v0.18.1. DNA methylation proportions were then calculated as the ratio of methylated reads to total reads.

RNA-seq data were summarized and normalized at the gene level using the R Bioconductor package *DESeq2* (v1.44.0) with the median-of-ratios normalization method implemented in *estimateSizeFactors()* [38, 39].

Promoter and linked enhancer regions were obtained from the GeneHancer “double elite” interaction track in the UCSC Genome Browser [40]. GeneHancer assigns enhancers to putative target genes using multiple lines of evidence, including eQTLs, Hi-C contacts, eRNA expression, transcription factor–gene coexpression, and genomic proximity, while the “double elite” subset retains only high-confidence regulatory interactions [41, 42].

Because genes may be linked to multiple enhancers located upstream, intragenically, or downstream, whereas promoter locations are uniquely defined, promoter and enhancer signals were summarized separately. We retained genes with both a promoter and at least one linked enhancer. Promoters were defined as 3-kb windows centered on the transcription start site (TSS; TSS ± 1,500 bp).

For each gene, we identified ATAC-seq peaks, histone-mark ChIP-seq peaks, and CpG sites overlapping either the promoter window or linked enhancer regions. We then constructed a gene-level regulatory score 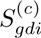 for gene *g*, data type *d*, sample *i*, and regulatory category *c* ∈ promoter, enhancer.

For ATAC-seq and ChIP-seq data, scores were computed as a width-weighted sum of peak intensities: 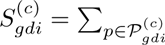 width*_p_,* height ∗ *p*, where *P*\**gdi*^(*c*)^ denotes the set of peaks assigned to (*g, d, i, c*).

For methylation data, the beta value at CpG site *j* was defined as 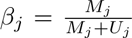, where *M_j_* and *U_j_* are the methylated and unmethylated read counts, respectively. Gene-level methylation scores were computed as coverage-weighted mean beta values across the *n* CpG sites assigned to 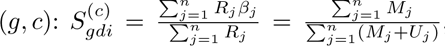, where *R_j_* = *M_j_* + *U_j_* denotes total coverage.

RNA-seq expression was normalized at the gene level using R bioconductor package *DESeq2* (v1.44.0) using the median-of-ratios method implemented in *estimateSizeFactors()* [38, 39]. After normalization, the log(1 + *p*) transformation and feature-wise standardization were applied so that the transformed data had a mean of 0 and a standard deviation of 1.

Overall, the IAV dataset included 54 final samples (27 subject in pre and post infection conditions) with sample-level characteristics age, ancestry (African American or European American) and infection status, and multiomic measurements for 18,187 genes with 13 data types (RNA-seq, and promoter and enhancer for each of ATAC-seq, DNA methylation, H3K27ac, H3K4me1, H3K4me3 and H3K27me3). Missing values occurred in promoter methylation, enhancer methylation and RNA-seq at rates of 5%, 14% and 9% respectively.

### Breast cancer dataset

The dataset used in our second study comes from the breast cancer collection in TCGA [10], accessed on March 12, 2026 via the XenaTools R package [43]. The TCGA BRCA cohort comprises multiomics molecular profiles for breast invasive carcinoma samples, and we analyzed three complementary data modalities: RNA sequencing–based gene expression, DNA methylation measured on the Illumina platform, and estimated copy number profiles (CNVs). We filtered samples based on the following clinically-relevant criteria: (1) female individuals, (2) cancer type was either infiltrating ductal/lobular carcinoma, (3) either pre or post menopausal (not peri), and (4) not missing survival information. The variables retained for analyses were:

1. Sample ID
2. Age at initial pathologic diagnosis - coded with three factors levels, *<*40, 40-60, *>*60.
3. Year of initial pathologic diagnosis - numeric.
4. Estrogen receptor status - positive or negative.
5. Progesterone receptor status - positive or negative.
6. HER2 receptor status - positive, equivocal or negative.
7. Pathologic stage - coded numerically as 1, 2 and 3.
8. Histological type - Ductal or Lobular.
9. Percent of positive lymph nodes - numeric between 0 and 1.
10. Progression free interval - binary indicator of progression event.
11. Progression free interval time - continuous, time variable for survival models. PFI was defined as in [15] as the interval from diagnosis to the first occurrence of disease progression, recurrence, distant metastasis, new primary tumor, or tumor-related death. Patients without these events were censored at their last follow-up or at death without disease.

For details on why these variables are of interest for breast cancer, see [44].

Gene expression data were generated using the Illumina HiSeq 2000 RNA-sequencing platform and represented as gene-level, log2(x + 1)-transformed RSEM-normalized counts. Copy number profiles were derived from whole-genome microarrays and processed using the GISTIC2 algorithm through the TCGA Firehose pipeline, yielding gene-level CNV measurements. DNA methylation data were generated using the Illumina Infinium HumanMethylation450 platform and represented as beta values ranging from 0 to 1. Gene expression and CNV features were mapped to HUGO gene identifiers using the UCSC Xena HUGO probeMap, while methylation probes were mapped using the UCSC Xena probeMap.

Gene coordinates were obtained from the Bioconductor package TxDb.Hsapiens.UCSC.hg38.knownGene [45], based on the UCSC hg38 reference genome. Entrez Gene identifiers were mapped to official HUGO gene symbols using the org.Hs.eg.db annotation database [46]. Promoter regions were defined for each gene as a 3 kb window centered on the annotated transcription start site (TSS). Methylation beta values were averaged within probes that fell in each gene’s promoter region.

We retained only samples with no missing data for the three omic measurements, leading to a dataset with 435 samples, each associated with 11 sample-level characteristics. In those samples there were 5,205 genes, including X chromosome genes, with no missing values in the omic measures.

## Declarations

### Ethics approval and consent to participate

This research was approved as project 2024-4041 by the Medical/Biomedical Committee of the Centre Intégré Universitaire de Santé et de Services Sociaux du Centre-Ouest-de-l’Île-de-Montréal.

### Consent for publication

Not applicable.

### Availability of data and materials

The raw data from the IAV dataset are available from the European Genome-phenome Archive (EGA), under accession numbers EGAD00001008422 and EGAD00001008359.

The TCGA breast cancer dataset analyzed during the current study were accessed via the Rpackage UCSCXenaTools version 1.7.0 [43] on March 12, 2026.

The code used to analyze the data is available in the MOLA repository, https://github.com/GreenwoodLab/MOLA.

### Competing interests

The authors declare that they have no competing interests.

### Funding

The authors acknowledge funding from the DNA-to-RNA (D2R) Foundational Projects (Cycle 1) grant: modeling individual differences in patterns of epigenetic regulation.

### Authors’ contributions

All authors conceived the experiments, S.B. conducted the experiments and analyzed the results. B.X. and K.O.K. preprocessed the data. S.B., C.G. J.D. and Q.Z. all reviewed the manuscript.

## Acknowledgments

We thank Dr. Patricia Tonin for her helpful suggestions regarding the selection of clinically-relevant variables and subgroups for our breast cancer study.

## Notes

https://github.com/GreenwoodLab/MOLA

### Competing Interest Statement

The authors have declared no competing interest.

