## Supplemental Figures for "MOLA: a novel topological data analysis framework for analyzing multiomic loops in precision medicine"

### Supplementary Figures

The following plots show gene set enrichment analysis (GSEA) results for GO [1], KEGG [2], and Reactome [3] pathways from our Influenza A virus epigenetic alteration analysis (Figure 2C). Enrichment analyses were performed using the R packages clusterProfiler and ReactomePA (see Methods for details). Each plot displays FDR-significant enriched pathways, with the normalized enrichment score (NES) on the x-axis. Point size represents the number of genes from the ($M = 500$) gene set assigned to each pathway, and point color indicates the enrichment p-value.

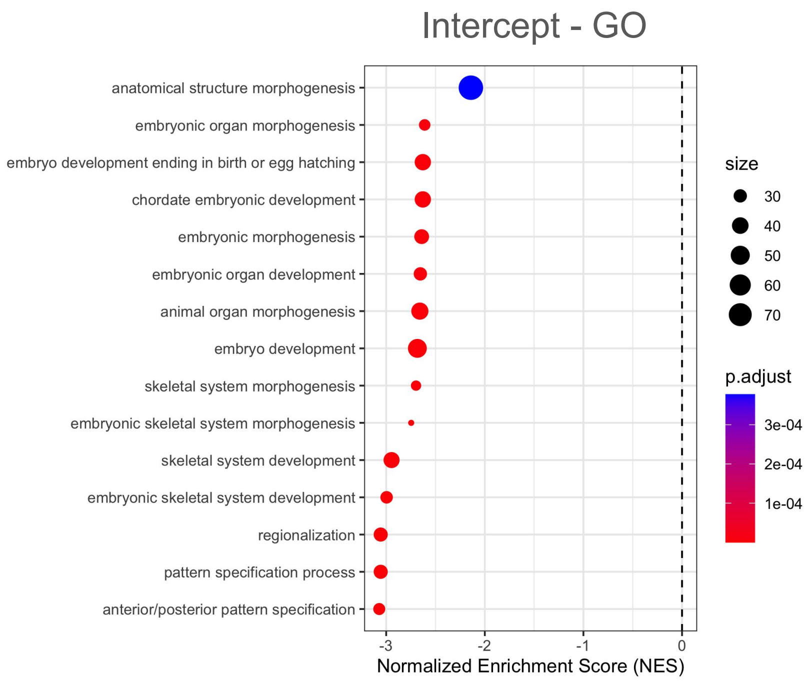


**Fig. S1**: Baseline (i.e., model intercept t-values) GO enrichment plot.

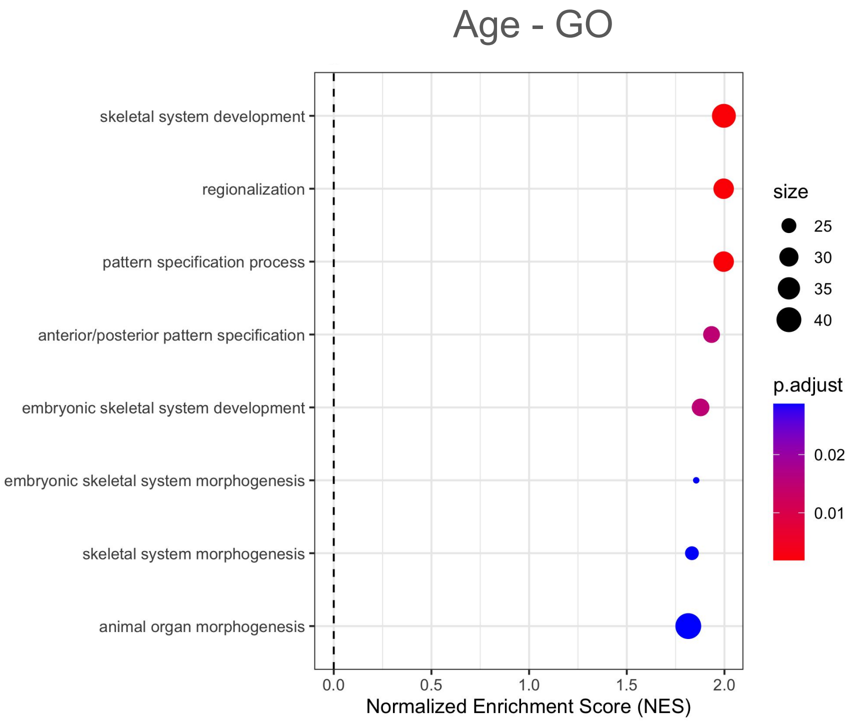


**Fig. S2**: Age GO enrichment plot.

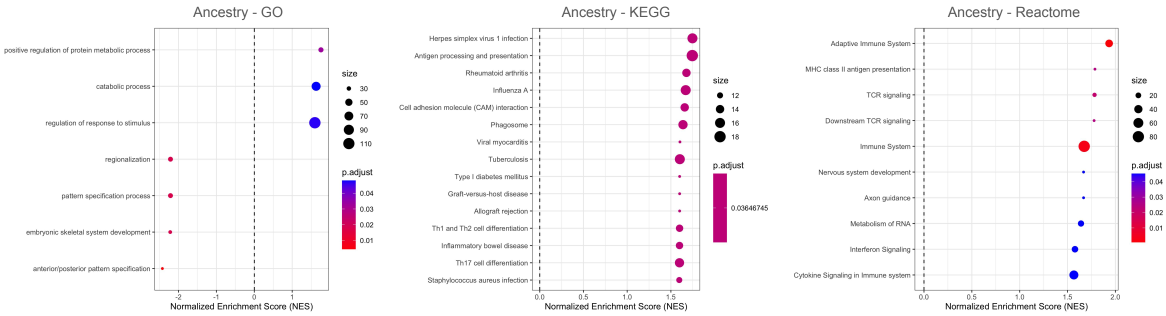


**Fig. S3**: Ancestry GO, KEGG & Reactome enrichment plots.

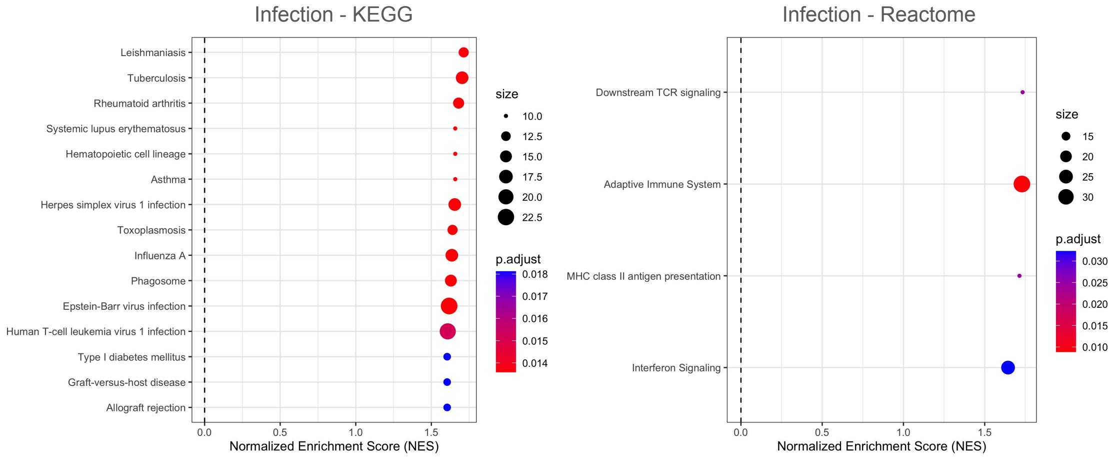


**Fig. S4**: Infection KEGG & Reactome enrichment plots.

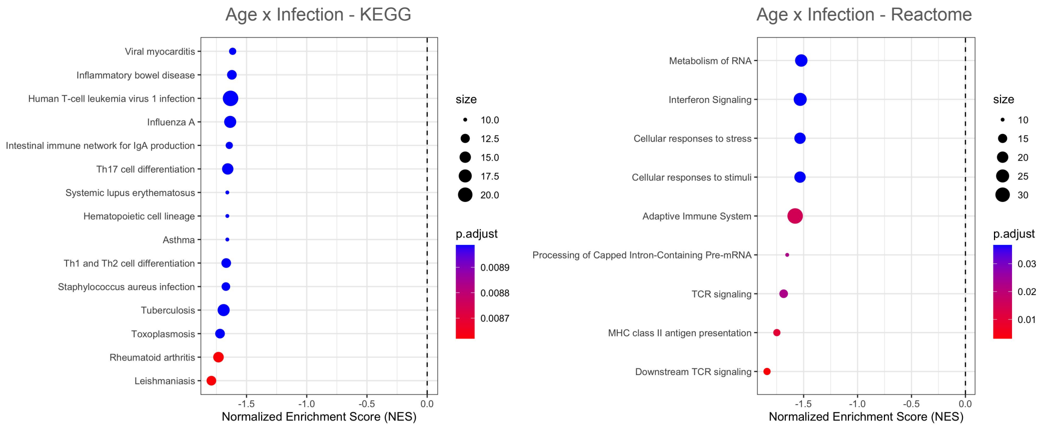


**Fig. S5**: Age x Infection interaction term KEGG & Reactome enrichment plots.

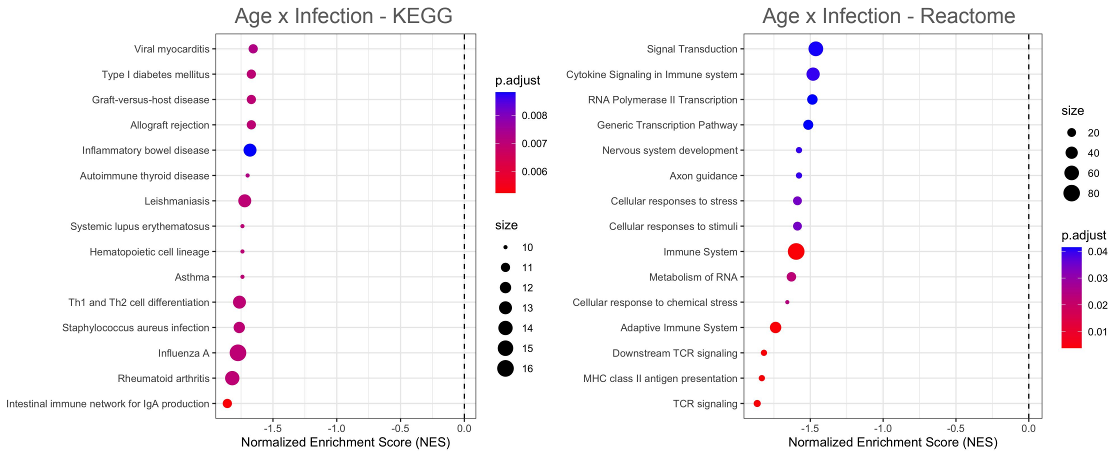


**Fig. S6**: Ancestry x Infection interaction term KEGG & Reactome enrichment plots.
